# What exactly is ‘*N*’ in cell culture and animal experiments?

**DOI:** 10.1101/183962

**Authors:** Stanley E. Lazic, Charlie J. Clarke-Williams, Marcus R. Munafò

## Abstract

Biologists establish the existence of experimental effects by applying treatments or interventions to biological entities or units, such as people, animals, slice preparations, or cells. When done appropriately, independent replication of the entity-intervention pair contributes to the sample size (N) and forms the basis of statistical inference. However, sometimes the appropriate entity-intervention pair may not be obvious, and the wrong choice can make an experiment worthless. We surveyed a random sample of published animal experiments from 2011 to 2016 where interventions were applied to parents but effects examined in the offspring, as regulatory authorities have provided clear guidelines on replication with such designs. We found that only 22% of studies (95% CI = 17% to 29%) replicated the correct entity-intervention pair and thus made valid statistical inferences. Approximately half of the studies (46%, 95% CI = 38% to 53%) had pseudoreplication while 32% (95% CI = 26% to 39%) provided insufficient information to make a judgement. Pseudoreplication artificially inflates the sample size, leading to more false positive results and inflating the apparent evidence supporting a scientific claim. It is hard for science to advance when so many experiments are poorly designed and analysed. We argue that distinguishing between biological units, experimental units, and observational units clarifies where replication should occur, describe the criteria for genuine replication, and provide guidelines for designing and analysing *in vitro, ex vivo,* and *in vivo* experiments.

## Introduction

Designing experiments is challenging: there are many options to consider, decisions to make, and trade-offs to weigh, and a single poor design choice can make an experiment worthless. We focus here on replication—a critical part of designing an experiment that is often misunderstood by experimenters.^1–7^ Misunderstandings inevitably lead to poor design choices, which in turn contribute to irreproducible or meaningless results.

Both statisticians^9–11^ and biologists^3,7,12–14^ agree on the importance of replication and distinguish between two types. The first is replication that increases the sample size (*N*) and thus contributes to testing an experimental hypothesis. It is called *true, genuine,* or *absolute* replication, and when these qualifiers are not used, *replication* is understood to mean this type. The second type is replication that does not increase the sample size and is called *pseudoreplication.^3^* An example will illustrate the difference: suppose a researcher hypothesises that male mice have heavier brains than female mice. He could (1) weigh the brain of one male and one female mouse five times, or (2) weigh the brain of five male and five female mice once. Both designs provide ten data points to calculate a p-value, but the p-value is meaningless for the first design because the hypothesis is about sex differences, and there is only one member of each sex. The multiple measurements on these two mice do not contribute to *N* and thus constitutes pseudoreplication. The wrong choice of replicate cannot be fixed by a clever statistical analysis after the experiment is completed; it needs to be planned at the design stage. Confusing pseudoreplication for genuine replication artificially inflates the sample size, thereby inflating the apparent evidence supporting a scientific claim and contributes to irreproducible results.

Unlike the above example, sometimes it is less clear what aspect of an experiment should be replicated to increase the sample size, and far too often the wrong aspect is chosen, leading to meaningless results^1–7^. Confusing pseudoreplication for genuine replication is an old problem; in 1929 Dunn discussed the distinction in *Physiological Reviews* and placed it first on his list of twelve admonishments to researchers.^1^ Papers and books warning of the genuine/pseudoreplication distinction have appeared regularly (four papers^15–18^ and a book chapter^19^ in the past year), but, paraphrasing Goodman’s comment on misinterpreting p-values, “these lessons appear to be either unread, ignored, not believed, or forgotten as each new wave of researchers is introduced to[…] research”.^20^

What do we hope to achieve with yet another discussion,? First, we argue that the frequent and common distinction made between “biological” and “technical” replication is unhelpful because they are inconsistently defined, do not capture the important characteristics of an experiment, and do not clarify what to replicate, and can therefore be used to justify an inappropriate design. We introduce instead the concept of biological units, experimental units, and observational units, and argue that they can clarify where replication needs to occur.^19^ Second, we provide detailed criteria for genuine replication that are applicable to all biological experiments and that researchers can use to design their experiments. We also remark on the assumptions that are made if a criterion is not met. Finally, we provide concrete examples to help translate the general principles and criteria into practice.

We begin by discussing the source of the problem—why is it hard to identify genuine replication? Next, we assess the extent of the problem by reviewing the literature on animal experiments where interventions were applied to parents but effects were examined in the offspring. We focused on these experiments because using the offspring as the sample size meets none of the criteria for genuine replication, and regulatory authorities have provided clear guidelines on replication with these experiments^21,22^. Finally, we describe the requirements for genuine replication and provide examples for *in vivo, ex vivo,* and *in vitro* experiments.

## Why is it hard to identify genuine replication?

One reason for the confusion between genuine and pseudoreplication stems from multiple levels of biological organisation. Recall the “hierarchy of life” from your first biology class, which starts with atoms at the bottom and ends with the biosphere at the top. Most biomedical research takes place in the middle of the hierarchy, between macromolecules (e.g. DNA, protein complexes) and organisms such as animals or patients. Multiple levels of biological organisation complicate experiments in two ways. First, properties at one level of biological organisation tend to be influenced by those above; for example, the properties or characteristics of an eye (organ) depend on the rat in which it is located (organism). We expect visual acuity to differ between old and young rats and between healthy and diseased rats, and it follows that two eyes from the same rat will tend to be more alike than eyes from two different rats. If the rats are similar, the variation within rats might be of a similar size to the variation between rats, but some additional rat-to-rat variation is expected. The same principle applies throughout the hierarchy; two cells in the same eye will tend to be more alike than cells between different eyes (within a rat), and two mitochondria within a cell will tend to be more alike than mitochondria between two different cells. Thus, if we are interested in visual acuity, “Rat” is a variable that predicts or affects visual acuity, before any experimental intervention is applied.

The second problem caused by multiple levels of biological organisation is that the research hypothesis, the experimental manipulations or interventions, and the measurements, could all be at different levels, making it unclear which level should be used to determine the sample size—is *N* the number of rats, eyes, cells, or mitochondria? Entities quickly multiply as we go lower in the hierarchy, each rat has two eyes, each eye has many cells, and each cell has many mitochondria. Are we free to choose what *N* refers to? Few problems arise when the hypothesis, manipulations/interventions, and measurements are restricted to one level of biological organisation, but many experiments span multiple levels.

In addition, technical hierarchies can make experiments more complex; for example, an *in vitro* experiment using a single cell line might be independently replicated on three separate days, with wells in a microtitre plate randomised to different treatment conditions, and measurements taken on several cells within each well. Is the sample size the number of days, wells, or cells? Or, does *N* = 1 because only one cell line was used?

While biological and technical hierarchies make identifying genuine replication a challenge, a contributing factor to the poor quality of many published studies is that biologists rarely receive formal training in experimental design and so cannot derive the answer from first principles. It is hard to think of another profession where a core competence is so infrequently taught to its members.

## What is the extent of the problem?

We searched PubMed for papers where experimental interventions were applied to parents but effects were examined in the offspring as these studies have an experimental intervention at one level (parents) and observations at the level below (offspring). The following search term was used: “prenatal exposure [TIAB] AND offspring [TIAB] AND (“2011/01”[DP]: “2016/09”[DP]) NOT review [PT]”, restricting the search to non-review articles between 2011 and September 2016. 500 abstracts were returned and from this list we randomly selected 200 abstracts to examine further. The inclusion criteria were: (1) English language paper, (2) multiparous mammals used in the experiments, and (3) paternal or maternal animals assigned to treatment conditions and the treatment applied to them, but the outcomes measured in the offspring. These criteria excluded studies on humans, nonhuman primates, and non-mammalian species. If a paper was deemed irrelevant (e.g. *in vitro* study) it was discarded and another randomly selected paper was included until we reached 200 papers that met the inclusion criteria. We chose 200 papers as that gives a target 95% confidence interval (CI) width of approximately 10%. Based on a previous study^23^ we expected that 80-90% of papers (160-180 out of 200) would mistake pseudoreplication for genuine replication, which gives a 95% CI width of between 9% and 12% (the CI width depends on the percentage of studies with pseudoreplication).

For each paper, we assessed whether *N* reflects genuine replication instead of pseudoreplication and classified papers into “Yes”, “No”, and “Unclear” categories. Unclear means that insufficient information was provided to make a Yes or No judgement. In addition, we recorded if the paper mentioned using randomisation and blinding, if the number of litters (or equivalently, the number of dams or paternal animals) and the number of offspring were reported, and if the experiment used a split-unit design. In a split-unit (also known as a split-plot) design, randomisation occurs at multiple levels in a hierarchy; for example, pregnant dams (and their unborn offspring) are randomised to treatment conditions, and when the offspring are born, they are randomised to another set of treatment conditions.

Since both blinding and randomisation can occur at several places in an experiment, as long as authors mentioned them once we assumed they were familiar with these methods and used them in all appropriate places (even if they were not mentioned again elsewhere). Papers with multiple animal experiments were classified as having used blinding or randomisation for all experiments if the authors mentioned these methods for at least one experiment. The data will be available at the data.bris Research Data Repository (http://data.bris.ac.uk/data/), doi: XXXXX, but have been attached as supplementary material for the initial submission.

The results of the literature survey were disappointing, but in line with previous findings^24–26^ (Fig. 1A). We found that only 22% of studies (95% CI = 17% to 29%) had genuine replication and thus made valid statistical inferences. Nearly half of the studies (46%, 95% CI = 38% to 53%) had pseudoreplication while 32% (95% CI = 26% to 39%) didn’t provide enough information to determine if *N* corresponds to genuine replication or pseudoreplication. We suspect that most studies in the Unclear category had pseudoreplication because if researchers were aware of this problem, they likely would have described how they avoided it.

**Figure 1:**
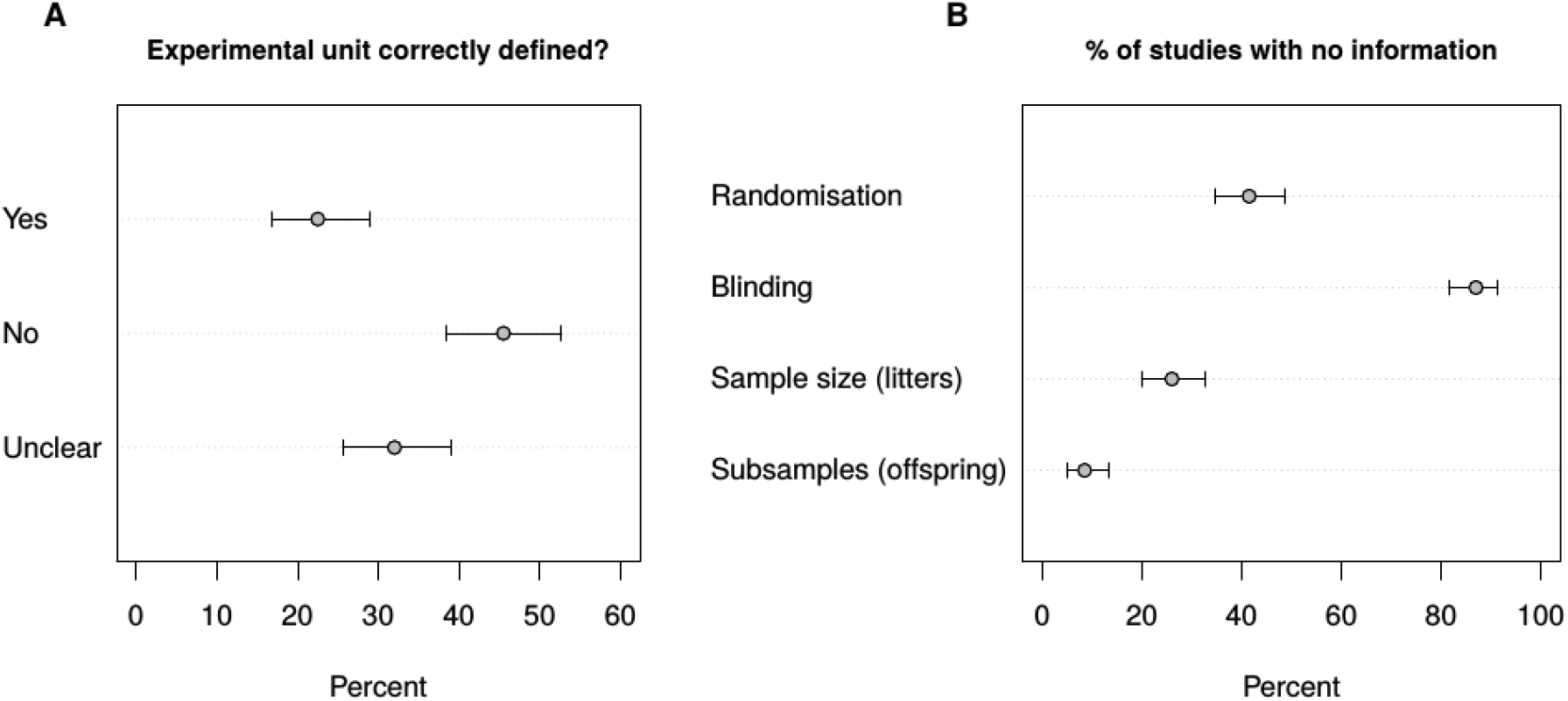
Results of the literature review. Nearly half of the studies (46%, 95% CI = 38% to 53%) confused pseudoreplication for genuine replication while 32% (95% CI = 26% to 39%) did not provide enough information to determine if the sample size was correct (A). Consistent with previous research, randomisation, blinding, and the sample size was not always reported (B). Error bars are 95% CI.

Furthermore, 42% of studies (95% CI = 35% to 49%) did not report using randomisation and 87% of studies (95% CI = 82% to 91%) did not report using blinding (Fig. 1B). In addition, 26% of studies (95% CI = 20% to 33%) did not report the sample size (number litters/dams/paternal animals) and 8.5% (95% CI = 5% to 13%) provided no information on the number of offspring. Half of the studies (48%; 95% CI = 41% to 56%) did not report the number of offspring for all experiments, or at least it was unclear how the total number of offspring was divided into subgroups for different experiments or readouts.

## Requirements for genuine replication

In experimental biology there is always a treatment or intervention applied to a biological entity such as a person, animal, or cell, and replication consists of multiple such entity-intervention pairs. For example, an intervention could be a drug given orally to a patient. Each patient-drug pair is considered one replication and provides independent information about the drug’s effect. The effect is usually relative and compared to a placebo or another drug. If the question is “what is the effect of the drug on patients” then replicating the patient-drug pair and patient-placebo pair enables the question to be addressed. Other aspects of this experiment could be replicated; for example, we could measure the outcome on each patient more than once and thus replicate the measurement procedure. Or, we could give the drug to the same person multiple times, and on each occasion record the outcome. We could also conduct the experiment multiple times, in different countries perhaps. Thus, decisions about what to replicate are unavoidable.

Regardless of the requirements for genuine replication (discussed below), there must also be replication relevant to the hypothesis being tested. For this it helps to define the scientific or *biological unit* (BU) of interest, which is the entity that we would like to test a hypothesis or draw a conclusion about, and common examples include people, animals, and cells. For example, many patients are required to conclude that a drug is better than a placebo because the hypothesis is about patients; we cannot give Jim the drug and Bob the placebo, take a daily measurement for several weeks, and then make conclusions about the drug’s efficacy in patients in general. At times, the BU may not be so apparent. Suppose we hypothesize that outbred mice are smarter than inbred mice. We cannot test this hypothesis with only one strain of outbred and one strain of inbred mice, even if we have many mice from each strain. There may be intelligence differences between these two strains that have nothing to do with their inbred/outbred status, much like there are differences between Bob and Jim, which cannot be disentangled from the drug effect. The biological unit is the strain (the hypothesis is about strains) and therefore we need multiple strains of both outbred and inbred mice.

Similar problems arise in cell culture studies. If we hypothesise that breast cancer cell lines proliferate at a faster rate than lung cancer cell lines, we need multiple breast and lung cell lines, as the hypothesis is about differences between the tissues of origin. If one breast and one lung cell line proliferate at different rates, we cannot attribute this to the tissue of origin as no two cell lines are expected to proliferate equally. Similarly, if we are interested in comparing proliferation rates between cancer cells with *p53* and *KRAS* mutations, then multiple cell lines with these mutations are required.

Having defined the biological unit in the experiment, the next entity to define is the *experimental unit* (EU), which is the entity that is randomly and independently assigned to the treatment conditions. Examples include a person, animal, litter, cage or holding pen, fish tank, culture dish, or well in a microtitre plate. In many experiments the biological and experimental unit might be the same entity, and the key point, which we return to, is that the sample size corresponds to the number of EUs.^10,11,19,27,28^

Finally, we can define the *observational unit* (OU) in an experiment, which is the entity on which measurements are made (Box 1). When all three units correspond to the same entity, the design and analysis of an experiment is simpler, but when the units refer to different entities, the intuition behind what constitutes *N* breaks down.

#### BOX 1 Types of units (adapted from Lazic^19^).

**Biological unit of interest (BU):** is the entity about which inferences are made. The purpose of an experiment is to test a hypothesis, estimate a property, or draw a conclusion about biological units.

**Experimental unit (EU):** the entity that is randomly and independently assigned to experimental conditions. The sample size (*N*) is equal to the number of EUs.

They may correspond to:

1. A biological unit of interest
2. Groups of biological units
3. Parts of a biological unit
4. A sequence of observations on a biological unit

In addition to random and independent assignment, for genuine replication:

1. The treatment should be applied independently to each EU, and
2. The EUs should not influence each other.

If these conditions do not hold, using a different unit as the EU may be preferable, usually a unit one level up in the biological or technical hierarchy.

**Observational unit (OU):** the entity on which measurements are taken, which may be different from the experimental and biological units of interest. Increasing the number of OUs does not increase the sample size.

Although the EU often corresponds to a biological unit of interest (Fig. 2A), it can also correspond to a collection or group of biological units, such as all mice in a litter if the treatment is applied to the pregnant dam (Fig 2B). In addition, the EU can correspond to parts of a biological unit; for example, individual eyes, patches of skin, or organotypic slice cultures from the same animal, as long as these parts can be randomly and independently assigned to different conditions (Fig 2C). Finally, an EU can correspond to a sequence of observations on a single biological unit. For example, the experiment is divided into time periods that are randomly assigned to different treatment conditions (e.g. on or off a drug), and a measurement is taken at each time period (Fig. 2D). This last design is infrequent in biological experiments but forms the basis for *N*-of-1 and cross-over designs. Below we use concrete examples to illustrate these points.

**Figure 2:**
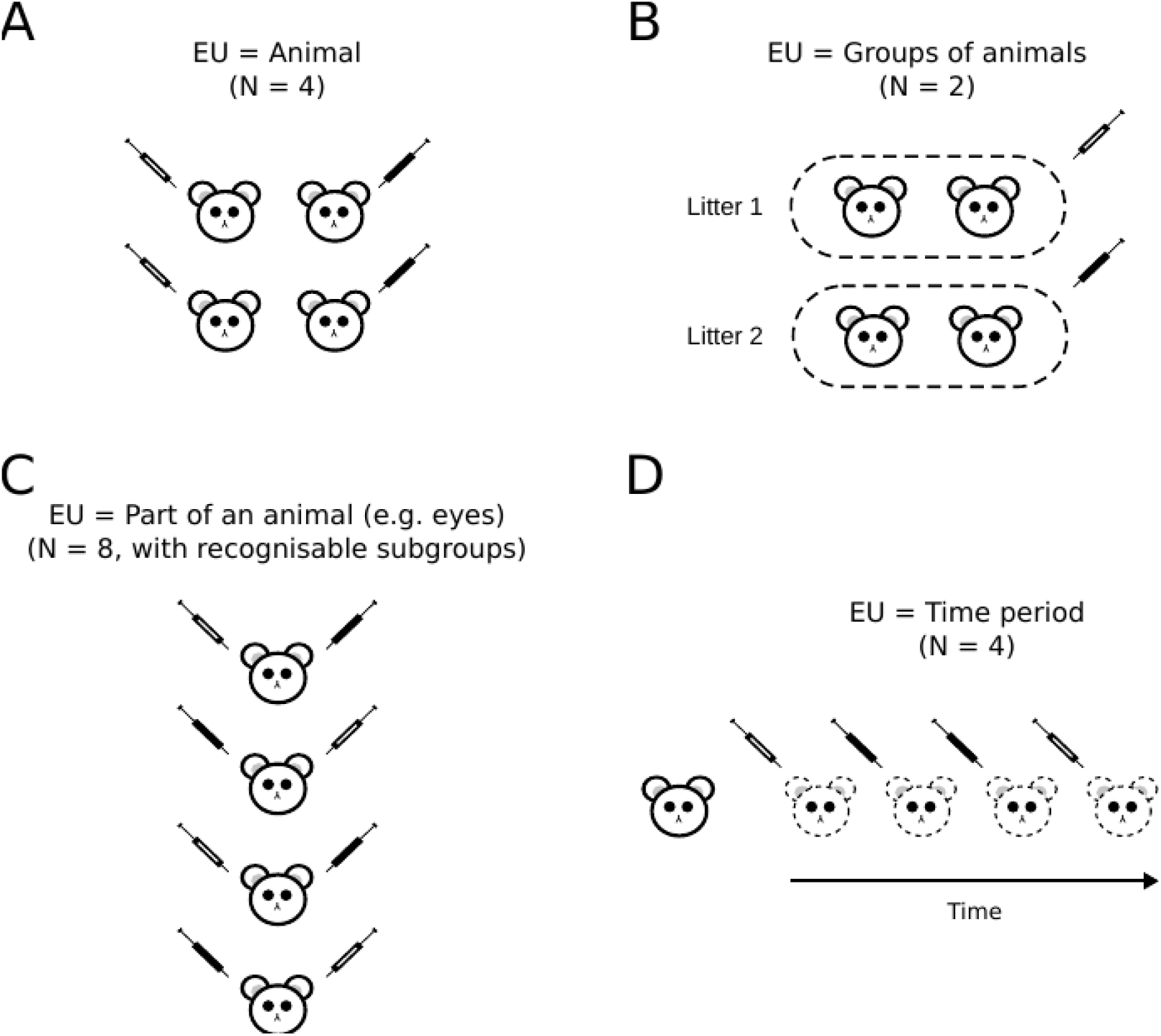
Types of experimental units (EU). An EU can correspond to a biological unit of interest such as an animal (A), groups of biological units, when they are randomised by group or when the treatment is applied group-wise (B), or parts of a biological unit, where each animal forms the “recognisable subgroup” (C). A less common example is when an experiment is divided into time periods that are randomly assigned to different treatment conditions and a measurement is taken during each time period. Note how the number of animals may differ from the sample size (N). The two experimental conditions are represented with syringes (white = control, black = treated). Adapted from Lazic.^19^

As mentioned previously, to increase the sample size we need to increase the number of experimental units (a useful phrase to remember is *“sample size is where you randomise”).^19^* But there are three conditions that must be met for an EU to be considered a genuine replicate (Fig. 3). First, EUs must be independently allocated to experimental conditions or interventions. Independence is required because p-value calculations assume independence. A p-value measures the discrepancy of the observed results to a theoretical null distribution. Since the null distribution is based on hypothetical alternative randomisations of the EUs to experimental conditions, we get a different null distribution (and p-values) if potential EUs are randomised independently or in batches or subgroups (e.g. animals in the same litter always end up in the same condition). A related concern is how these subgroups are formed. Are they natural groupings of biological units such as animals in a litter, or all cells in a tissue sample? If so, then we expect the potential EUs within a subgroup to be more alike on relevant variables than EUs between subgroups, as discussed earlier. When the pool of EUs to be randomised have such “recognisable subgroups”, which are typically defined by biological factors such as sex, litter, age, and so on, it makes sense to ensure that the subgroups are balanced across the treatment conditions. A balanced design can be acheived by randomising within each subgroup separately (Fig. 3, right side).

The second requirement is that the experimental intervention must be applied independently to each EU. Independence is needed here because we cannot exactly replicate the application of a treatment to all EUs—known as *treatment error.* For example, intraperitoneal injections are never in exactly the same location and the same amount, but when given independently to each rodent these treatment errors average out across the experimental groups. If we accidentally give a rodent twice the required dose, only this animal will be affected. But if instead we apply the treatment to groups of animals simultaneously, then all animals will be affected the same way (the treatment errors are correlated). For example, if we accidentally give twice the concentration of a drug in the drinking water, then all animals in the cage will be affected. In many experiments the treatment error will be small compared with the size of the treatment effects we are trying to detect, and barring any experimental mistakes, the treatment error may be negligible. But this is an assumption, and one that is usually untestable. A related point is that the treatments should not spill over or affect adjacent experimental units, especially those in another condition. For example, if one side of the brain is infused with a growth factor and the other side serves as a control, we assume that the growth factor does not diffuse to the opposite hemisphere. If we cannot be sure that this assumption holds, and especially if we are unable to verify it, then it may be better to use different animals for the treated and control conditions.

The third requirement is that EUs must not influence each other, especially on the measured outcomes variables. For example, Kalbassi recently found that the behaviour of wild-type mice was altered when they were housed with their Neuroligin-3 knock-out littermates (a model of autism).^29^ Here, EUs in one condition are affecting EUs in another. But EUs within the same experimental condition must not influence each other on the relevant outcomes either. Consider a group of students taking a statistics test that are in the same experimental condition and meet all of the above requirements for genuine replication. If the examiner leaves the room and the students collaborate, we expect the variability of the test scores to decrease. The students copy each others’ answers and their responses become more alike (not independent). If we use students as the EU, the variability amongst the test scores is too low, leading to artificially precise estimates, which usually translates into p-values that are too small. The mean test score of all students is the only useful piece of information. (If weaker students tend to copy the correct answers from stronger students, then the mean test score will be biased, but if only the variability is affected, then the group mean can still be used in an analysis).

We expect animals in a cage to influence each other on many relevant variables, from behaviour to microbiomes (and hence anything influenced by an animal’s microbiome). Even if animals meet the first two criteria for genuine replication, mutual influence of animals in the same cage may render them unsuitable to be an EU. The solution is to house animals one per cage, or, if this is undesirable for ethical or experimental reasons, housing animals two per cage maximises the number of cages (which are now the EUs) for a fixed number of animals. The same idea applies to cells in a well or tissue, if there is mutual influence, or suspected influence, then *N* cannot refer to the number of cells. In the following subsections we show how these ideas apply to different types of experiments.

**Figure 3:**
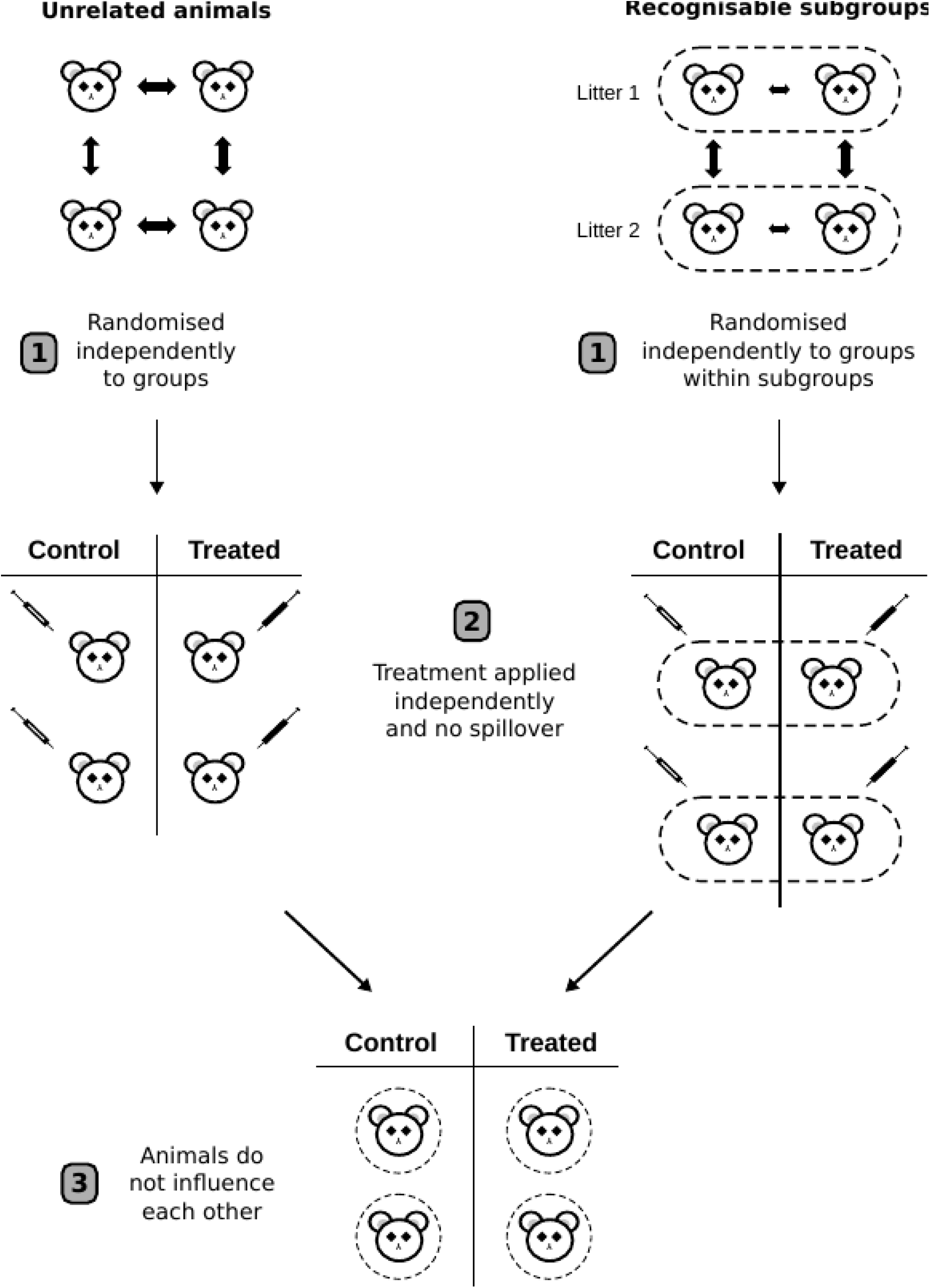
Three criteria for genuine replication. (1) Experimental units (animals) must be independently randomised to the treatment conditions. Double arrows indicate variability between EUs, and is constant when the EUs are homogeneous. This variability between the EUs is used to test for differences between treatments. If there are recognisable subgroups such as litters or sex, randomisation can be done within each subgroup. Variability will tend to be less between EUs in a subgroup (smaller arrows) than between subgroups. (2) The treatment must be independently applied to each EU and must not affect or spillover to adjacent EUs. The syringes represent the application of an intervention to the EUs (white = control, black = treated). (3) The EUs must not influence each other (dashed circles represent animals isolated from influencing each other),especially on the outcomes of interest.

## Animal experiments

If treatments are randomly and independently applied to an entity other than the individual animal, then the sample size is not the number of animals. The studies in the literature review applied treatments to pregnant females and observed the effects in the offspring. None of the criteria are met for using the offspring as genuine replicates: animals in the same litter are randomised together, the treatment is applied to all animals simultaneously, and it is unreasonable to assume that animals in a womb do not influence each other (e.g. competing for maternal resources). In these experiments the experimental units (pregnant dams or occasionally the fathers) do not correspond to the biological unit of interest (offspring)^23^, and regulatory authorities do not allow the offspring to be used as independent samples for toxicology studies using this design.^21,22^

What about a similar experiment where, after offspring are born, they are randomised by litter to treatment groups, so that all animals in a litter end up in the same treatment group. Now the treatment can be applied independently to each animal (criteria 2) and animals can be prevented from influencing each other (criteria 3). But since the animals are not independently randomised (although they could have been), the first criterion is not met and the litter is the EU. This experiment may be less misleading if offspring are treated as the EU since two of the three criteria are met, but litter is still the appropriate EU. Another problem with this design is that the variable *litter* is nested under the variable *treatment group*,which is a less powerful design compared with crossing litters and treatment groups.^19^ In the crossed arrangement (Fig. 3, right side) the individual animals are randomised to the treatment condition, so *N* is the number of animals. This is a powerful design even if there are large differences between litters because the litter-to-litter variation can be cleanly estimated and removed (assuming ‘litter’ is included as a variable in the analysis). A nested arrangement is rarely a good idea because the BU and EU do not coincide, it is harder to analyse, and is less powerful than the crossed design. The same idea extends to more than two animals per litter and more experimental groups.

In some experiments unrelated animals may be (1) randomised to treatment groups and then housed together by treatment group, or (2) randomised to cages and then cages are randomised to treatment groups. In both cases animals, and not cages, can be considered the EU because the animals are independently assigned to the treatment groups (assuming that the second and third criteria are also met).

In other experiments unrelated animals may be randomised to treatment groups and the treatment applied cage-wise in the drinking water or to individual animals by gavage. When a treatment is applied cage-wise, the cages are the experimental units because the treatment is applied simultaneously to all animals in a cage (second criterion not met), whereas when the treatment is applied independently to each animal, the animal is the EU. For many experiments the difference between these two designs may be negligible if the treatment error is low, but nevertheless, it is important to know the assumptions made.

Non-human primate experiments tend to have small sample sizes for both cost and ethical reasons, but the requirements for genuine replication remain the same. Suppose an experiment records from a single neuron while pictures of a happy monkey or a sad monkey are shown to the subject. One hundred pictures are shown in random order, and for each trial the firing rate of the neuron is measured. We find that the neuron fires at a faster rate in the happy monkey condition, and the sample size is the 100 trials (this design resembles Fig 2D). This seems to be an easy way of obtaining a large sample size with only one subject and only one neuron, but the catch is that the results apply only to this subject and to this neuron, and little can be said about what might be seen in other subjects or other neurons. A statistical test would be valid, but the hypothesis tested is uninteresting. One may argue that this subject is representative of others, but this is a non-statistical generalisation, and the smallness of the p-value does not provide more evidence about what might happen in other subjects (showing that a drug works for Jim (p < 0.001) does not provide strong evidence that it works for Bob, or anyone else).

Suppose this experiment includes a second subject, run under identical conditions, with similar results. There are now 200 data points, 100 to test for an effect *within each subject*, but still only *N* = 2 to test for an affect across subjects (showing that a drug works for Jim (p < 0.001) and that it also works for Bob (p < 0.00001) provides little evidence about what might be seen in others). There are two hypotheses here, which must be reflected in the analysis and interpretation. Strong conclusions cannot be made about subjects in general from these two subjects.

In another variation, suppose that we have only one subject, but can record from ten neurons simultaneously. *N* is not increased ten fold because showing a picture is the application of the treatment, and it is applied to the subject and to all neurons simultaneously (2nd criterion is not met, much like applying a drug to all the cells in a well). Furthermore, we cannot rule out that the neurons are not influencing each other—they may be connected with gap junctions or have synaptic connections for example (3rd criterion is not met).

## Slice preparations and histological samples

In some experiments, animals are randomised to treatment conditions and an intervention is applied to the animals. Then, an organ or body part is examined, usually postmortem. The experimental unit is the animal and the sample size is the number of animals. Multiple histological sections, neurons per section, spines per neuron, and so on, are all subsamples as they have been randomised together (1st criterion is not met), the treatment is applied simultaneously (2nd criterion is not met), and they may influence each other between the treatment application and when the tissue is fixed (3rd criterion is not met).

In other experiments, a body part is first removed from the animal, the treatment applied, and then observations are made. If there are multiple body parts per animal such as the left and right hippocampus, two kidneys, or several blood samples, each of these can be randomised to different treatment conditions. The situation is identical to the right side of Figure 3, but instead of litters, the subgroups are the animals and the body parts are randomised to treatment groups within animals. Here, *N* is the number of body parts, but it is still useful to have multiple animals as this can establish the robustness of the effect, allow one to assess how the effect varies across animals, and makes generalisation to other animals possible.

Some *ex vivo* experiments are similar to the nonhuman primate example above, where, for example, an organotypic hippocampal culture has time periods randomised to the presence or absence of a serotonin receptor antagonist, and at each period electrophysiological recordings are made. Here we substitute the hippocampal culture for the subject, and the presence/absence of the antagonist for the pictures, and all previous points apply. In other *ex vivo* experiments, the treatment is applied to the animal, and multiple slice preparations are derived from each animal. Here, *N* is the number of animals not the number of slice cultures.

## Cell culture experiments

In cell culture experiments cells are often both the observational and biological units of interest, but rarely the experimental unit. Suppose an aliquot of cells is thawed and the cell suspension is pipetted into different wells of a microtitre plate. Cells are randomised to wells, and then wells to treatments, so the first criterion is met. But treatments are applied simultaneously to all cells in a well, not independently to each cell, so the second criterion is not met. In addition, it is unreasonable to assume that cells in a well have no influence on each other; they form cell-to-cell connections, release signalling molecules, and compete for the same nutrients in the media. Hence, the third criterion is not met for using cells as the EU. Thus, a well, culture dish, or another plastic container is a suitable EU for cell culture experiments.

But *in vitro* experiments are often finicky; the results depend on the unique conditions that vary each time the experiment is run. The experimental material (e.g. a cell line) is often artificially homogeneous and the conditions under which the experiment is run are so narrowly defined that it is hard to know what will happen if the experiment is run a second time. For this reason, *in vitro* experiments are usually repeated on multiple days, and the number of wells, aliquots, or culture dishes within a day are treated as subsamples. The aim is to establish that the phenomenon is robust enough to survive multiple replications of the entire experimental run or protocol in a highly artificial system. This situation has parallels to the nonhuman primate example above. We could do the experiment on one day and use the wells as the EU, and have a large sample size, but then we cannot comment on the generalisability of the results. If the experimental system is sensitive to the many details of how it is carried out, then repeating the whole procedure on multiple days provides further information in a way that using more wells on a single day does not. It provides an estimate of the consistency of the effects across the different experimental runs (days). The multiple wells on each day are then treated as subsamples and do not contribute to *N* (for example, by averaging values across wells in the same condition on each day). This is a scientific judgement about the relevant unit that we would like to make inferences about, and although opinions may differ, using more stringent criteria makes the results more believable.

## Analysis of hierarchical data

The analysis of hierarchical data is a large topic and beyond the scope of this paper, but a simple solution to the problem of pseudoreplication is to reduce the data to a single value for each experimental unit. Reducing the data by calculating the mean of multiple measurements on each EU will often be appropriate, but other numeric summaries such as the slope or area under the curve may capture a feature of interest better^30^. These summary-measure or derived-variable analyses allow standard statistical tests to be used, and will often give the same answer as more complex hierarchical models.

## Conclusion

There are few ways to conduct an experiment well, but many ways to conduct it poorly. Without identifying the correct experimental unit and having replication in the right place, experiments will likely be of little value or will test an uninteresting hypothesis. Other aspects of experimental design and how to analyse data from the above examples are beyond the scope of this paper, but are discussed in detail elsewhere.^19^ We hope this presentation provides researchers with the necessary principles to get the most value out of their experiments.

## Acknowledgements

MRM is a member of the UK Centre for Tobacco and Alcohol Studies, a UK Clinical Research Council Public Health Research: Centre of Excellence. Funding from British Heart Foundation, Cancer Research UK, Economic and Social Research Council, Medical Research Council, and the National Institute for Health Research, under the auspices of the UK Clinical Research Collaboration, is gratefully acknowledged. Support from the Medical Research Council (MC UU 12013/6) is also gratefully acknowledged.

## Author contributions

CJC-W and MRM performed the literature search. CJC-W and SEL evaluated the manuscripts. SEL performed the power analysis and analysed the data. All authors contributed to planning the study and writing the manuscript.

## References

1. Dunn H. L. Application of statistical methods in physiology. Physiological Reviews 9, 275–398 (1929).

2. Haseman J. K. & Hogan M. D. Selection of the experimental unit in teratology studies. Teratology 12, 165–171 (1975).

3. Hurlbert S. H. Pseudoreplication and the design of ecological field experiments. Ecol Monogr 54, 187–211 (1984).

4. Prosser J. I. Replicate or lie. Environ Microbiol 12, 1806–1810 (2010).

5. Ramirez, C. C., Fuentes-Contreras, E., Rodriguez, L. C. & Niemeyer, H. M. Pseudoreplication and its frequency in olfactometric laboratory studies. Journal of Chemical Ecology 26, 1423–1431 (2000).

6. Morrison D. A. & Morris E. C. Pseudoreplication in experimental designs for the manipulation of seed germination treatments. Austral Ecology 25, 292–296 (2000).

7. Lazic S. E. The problem of pseudoreplication in neuroscientific studies: Is it affecting your analysis? BMC Neurosci 11, 5 (2010).

8. Altman D. G. & Bland J. M. Statistics notes. units of analysis. BMJ 314, 1874 (1997).

9. Box, G. E. P., Hunter, J. S. & Hunter, W. G. Statistics for experimenters: Design, innovation, and discovery. (Wiley, 2005).

10. Casella G. Statistical Design. (Springer, 2008).

11. Mead, R., Gilmour, S. G. & Mead, A. Statistical Principles for the Design of Experiments: Applications to Real Experiments. (Cambridge University Press, 2012).

12. Landis S. C. et al. A call for transparent reporting to optimize the predictive value of preclinical research. Nature 490, 187–191 (2012).

13. Vaux, D. L., Fidler, F. & Cumming, G. Replicates and repeats-what is the difference and is it significant? A brief discussion of statistics and experimental design. EMBO Rep 13, 291–296 (2012).

14. Vaux D. L. Research methods: Know when your numbers are significant. Nature 492, 180–181 (2012).

15. Mundry R. & Oelze V. M. Who is who matters-The effects of pseudoreplication in stable isotope analyses. Am. J. Primatol. 78, 1017–1030 (2016).

16. Forstmeier, W., Wagenmakers, E. J. & Parker, T. H. Detecting and avoiding likely false-positive findings - a practical guide. Biol Rev Camb Philos Soc (2016).

17. Colegrave N. & Ruxton G. D. Statistical model specification and power: recommendations on the use of test-qualified pooling in analysis of experimental data. Proc. Biol. Sci. 284, (2017).

18. Tincani F. H. et al. Pseudoreplication and the usage of biomarkers in ecotoxicological bioassays. Environ. Toxicol. Chem. (in press), (2017).

19. Lazic S. E. Experimental Design for Laboratory Biologists: Maximising Information and Improving Reproducibility. (Cambridge University Press, 2016).

20. Goodman S. A dirty dozen: Twelve p-value misconceptions. Semin Hematol 45, 135–140 (2008).

21. International Conference on Harmonisation. Detection of toxicity to reproduction for medicinal products and toxicity to male fertility. S5(R2) (1993).

22. OECD. Guideline for the testing of chemicals: Developmental neurotoxicity study. 1–26 (2007).

23. Lazic S. E. & Essioux L. Improving basic and translational science by accounting for litter-to-litter variation in animal models. BMCNeurosci 14, 37 (2013).

24. Bebarta, V., Luyten, D. & Heard, K. Emergency medicine animal research: Does use of randomization and blinding affect the results? Acad Emerg Med 10, 684–687 (2003).

25. Macleod M. R. et al. Evidence for the efficacy of NXY-059 in experimental focal cerebral ischaemia is confounded by study quality. Stroke 39, 2824–2829 (2008).

26. Kilkenny C. et al. Survey of the quality of experimental design, statistical analysis and reporting of research using animals. PLoS One 4, e7824 (2009).

27. Festing M. F. W. Principles: The need for better experimental design. Trends Pharmacol Sci 24, 341–345 (2003).

28. Hinkelmann K. & Kempthorne O. Design and Analysis of Experiments, Volume 1: Introduction to Experimental Design. (Wiley, 2008).

29. Kalbassi, S., Bachmann, S. O., Cross, E., Roberton, V. H. & Baudouin, S. J. Male and female mice lacking neuroligin-3 modify the behavior of their wild-type littermates. eNeuro 4, (2017).

30. Matthews, J. N., Altman, D. G., Campbell, M. J. & Royston, P. Analysis of serial measurements in medical research. BMJ 300, 230–235 (1990).

